# Stroke disconnectome decodes reading networks

**DOI:** 10.1101/2022.03.20.485029

**Authors:** Stephanie J. Forkel, Loïc Labache, Parashkev Nachev, Michel Thiebaut de Schotten, Isabelle Hesling

**Affiliations:** Brain Connectivity and Behaviour Laboratory, Sorbonne Universities, Paris, France; Donders Centre for Cognition, Radboud University, Thomas van Aquinostraat 4, 6525 GD Nijmegen, the Netherlands; Centre for Neuroimaging Sciences, Department of Neuroimaging, Institute of Psychiatry, Psychology and Neuroscience, King’s College London, London, UK; Departments of Neurosurgery, Technical University of Munich School of Medicine, Munich, Germany; Department of Psychology, Yale University, New Haven, CT 06511, USA; UCL Queen Square Institute of Neurology, University College London, Queen Square, London, WC1N 3GB, UK; Groupe d’Imagerie Neurofonctionnelle, Institut des Maladies Neurodégénératives-UMR 5293, CNRS, CEA University of Bordeaux, Bordeaux, France

**Keywords:** *Language*, *disconnection*, *reading*, *stroke*, fMRI, *Exner*, VOF

## Abstract

Cognitive functional neuroimaging has been around for over 30 years and has shed light on the brain areas relevant for reading. However, new methodological developments enable mapping the interaction between functional imaging and the underlying white matter networks. In this study, we used such a novel method, called the disconnectome, to decode the reading circuitry in the brain. We used the resulting disconnection patterns to predict the typical lesion that would lead to reading deficits after brain damage. Our results suggest that white matter connections critical for reading include fronto-parietal U-shaped fibres and the vertical occipital fasciculus (VOF). The lesion most predictive of a reading deficit would impinge on the left temporal, occipital, and inferior parietal gyri. This novel framework can systematically be applied to bridge the gap between the neuropathology of language and cognitive neuroscience.

## Introduction

*This discovery of fire [is] probably the greatest ever made by man, excepting language [*…] (Darwin, 1871).

The invention of the written language is a pivotal hallmark in human history, and mastering reading and writing (i.e. literacy) requires many years. Written language is critical to society and spans from passing information between individuals to written rules that govern society. In today’s world, literacy abilities are indispensable to learning and navigating our social and professional environments.

Reading and writing require the interplay of several processes, including abstraction. For alphabetic languages (sound-based, e.g. English), when we write, we graphically encode (grapheme) linguistic utterances (phoneme) with the aim of a reader being able to decode and reconstruct the meaning during reading accurately. For this information transaction to succeed, one must understand the language associated with the symbols to comprehend the text.

Seminal work on the functional brain activations of proficiency in literacy has shown that reading selectively activates bilateral temporo-occipital areas, including the left visual word form area and right occipital areas (Dehaene et al., 2010; Murphy et al., 2019). However, reading proficiency modulates a more comprehensive left hemisphere language network, suggesting that literacy acquisition is tapping into the extended language circuitry in the brain. This network includes the left temporal cortex, premotor, and supplementary motor area (SMA) (Dehaene et al., 2010). Literacy-related activations also extended to the planum temporale suggesting a top-down modulation of orthography from spoken inputs. Although several studies have offered evidence that the left PT is involved in speech production and perception, some studies have also enlightened that the left PT is also recruited during word reading (Buchsbaum et al., 2005; Hesling et al., 2019). These changes occur regardless of when literacy was acquired (adulthood vs childhood) (Dehaene et al., 2010). The inferior frontal gyrus, harbouring ‘Broca’s area’, has been associated with various linguistic processes, including phonological segmentation, syntactic processing, and unification (Burton et al., 2000; Friederici, 2002; Hagoort, 2005). Common denominators to these processes are the aspects of segmenting and linking different types of linguistic information. While reading single words, in our study, highly automated words (i.e. days of the week) are unlikely to engage complex semantic and syntactic processing. Still, single-word reading requires linking phonemic sequences with motor gestures (Flinker et al., 2015) and verbal working memory, which may also recruit the inferior frontal gyrus (Jobard et al., 2003).

Literacy, therefore, requires a network of language-processing, auditory, and visual regions to interact. These relevant regions are distant from each other and, as such, rely on a network of connections. Classical language and auditory areas in the frontal and temporal lobes are connected by the arcuate fasciculus, the most prominent pathway for language functions (for review, see (Forkel et al., 2021). The fronto-temporal branch, also known as the long segment of the arcuate fasciculus, has also been shown to be relevant for learning new words via the phonological route (López-Barroso et al., 2013). The long segment is also associated with reading skills in children (Yeatman et al., 2011), while the parietal-temporal branch, also known as the posterior segment, has been associated with reading skills in adults (Thiebaut de Schotten et al., 2014). Another network that connects occipital-temporal-frontal cortices includes the inferior longitudinal fasciculus (ILF, occipito-temporal connection) (Catani et al., 2003), inferior fronto-occipital fasciculus (IFOF, occipital-frontal connection) (Forkel et al., 2014), and the vertical occipital fasciculus (e.g. VOF, intraoccipital connection) (Forkel et al., 2015; Sihvonen et al., 2021; Vergani et al., 2014; Yeatman et al., 2014). Longitudinal studies in children acquiring literacy revealed the left IFOF (Vanderauwera et al., 2018) and ILF (Yeatman et al., 2012) as mediators of orthographic (visual) but not phonological (auditory) processes. The VOF has been suggested as the link between the dorsal and ventral visual streams and to contribute to reading through the integration of eye movement with word recognition (anterior branch) and communication with visual processes (posterior branch) (Yeatman et al., 2014). In addition to the intrahemispheric communication between dorsal and ventral networks, information integration between both hemispheres (i.e. interhemispheric interaction) is critical to literacy. Reading relies on visual inputs from both eyes that must be synthesised in the brain. Accordingly, the posterior corpus callosum connects the parietal and occipital lobes to mediate this integration (Carreiras et al., 2009). While all these studies sporadically highlight pieces of brain networks that orchestrate the language, visual, and auditory processes necessary for reading acquisition, their relationship with the networks whose damage will contribute critically to reading impairment is not unequivocal.

Clinical neuroanatomical studies in patients who temporarily or chronically lost the ability to read after brain lesions only partially overlap with the networks related to literacy acquisition.

For instance, phonological dyslexia, an acquired dyslexia characterised primarily by difficulties in reading pronounceable nonwords, is usually caused by sizeable left hemisphere perisylvian lesions in the middle cerebral artery territory (Luzzatti et al., 2001). Surface dyslexia, the difficulty with whole word recognition and spelling, has been associated with lesions to the anterior lateral temporal lobe (Patterson and Hodges, 1992). While letter-by-letter dyslexia, characterised by prolonged reading and a linear word length effect, presents after lesions to the occipital and inferior temporal regions (Cohen et al., 2004; Fiset et al., 2006). Pure alexia, defined as pure word blindness by Dejerine (1892), has been initially characterised by lesions to the angular gyrus. However, this view has been challenged, and current models suggest an occlusion of the left posterior cerebral artery subserving parietal and occipital-temporal regions to cause the symptom (Cohen et al., 2000; Dehaene and Cohen, 2011; Philipose et al., 2007; Starrfelt and Shallice, 2014). Hence the relationship between the areas functionally activated when reading, the network required for literacy acquisition, and the critical network for reading disorders is missing.

A potential way to fill this gap is to take advantage of recent developments in functional-structural connectivity coupling that provides methods to decode functional MRI activations based on the pattern of brain disconnection derived from stroke – the disconnectome (Thiebaut de Schotten et al., 2020). This method reveals the white matter circuitry connecting activated areas and typical lesions that should interrupt this circuitry, thereby identifying the critical white matter circuitry for a specific cognitive function - in this case, reading. The disconnectome has never been applied in full to the reading circuitry. Therefore, in the present study, we reanalysed data (Hesling et al., 2019) in light of the disconnectome to reveal the reading circuitry and typical lesions that should impair its functioning.

## Methods

A methodological overview is provided in Figure 1. Briefly, we used previously published data (grey boxes) for a functional reading task (Hesling et al., 2019) and the disconnectome maps (Thiebaut de Schotten et al., 2020) to calculate the group-level statistical map for reading and use the disconnectome to decode the critical white matter network for reading.

**Figure 1.**
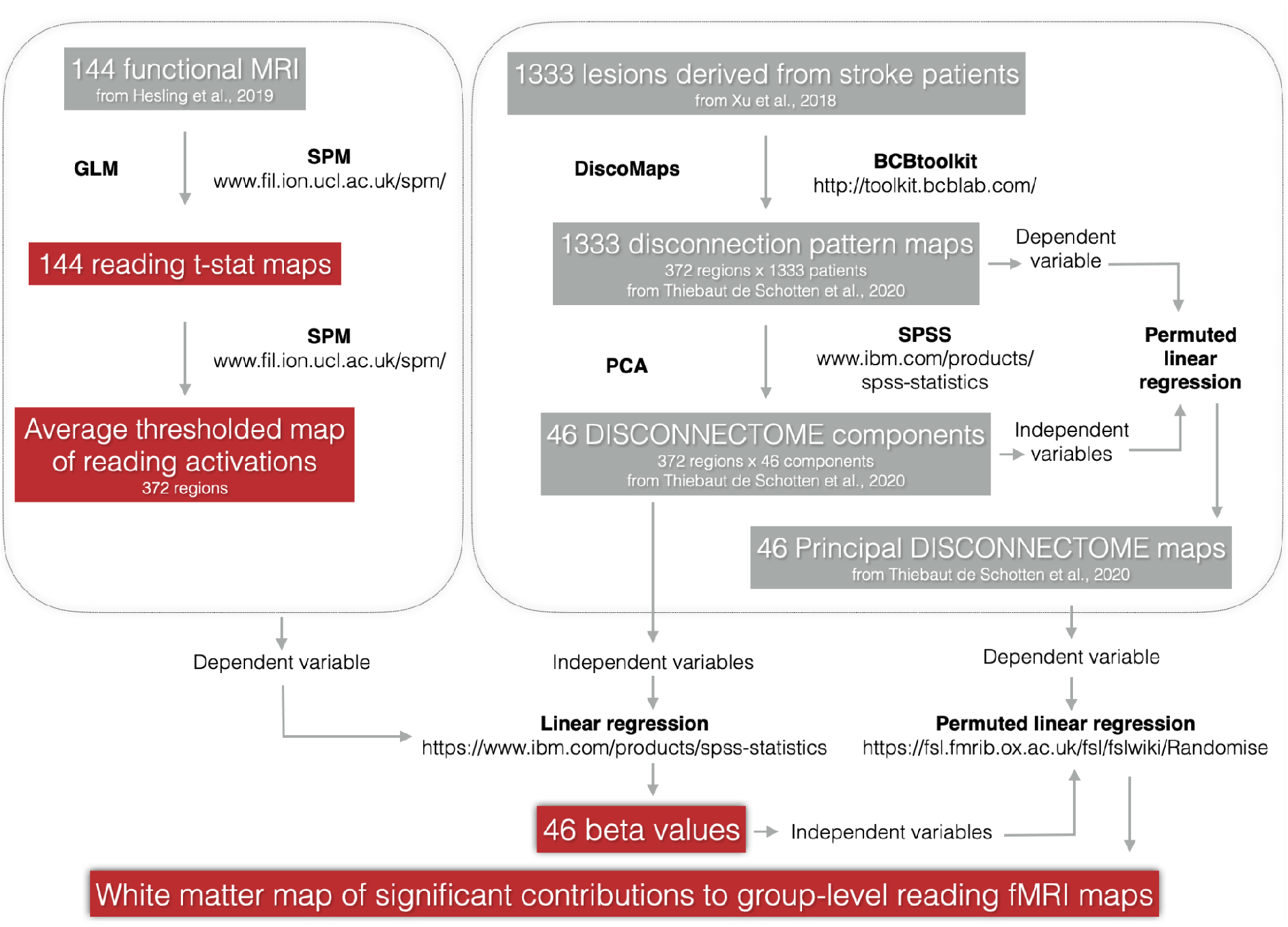
Schematic overview of the processing steps implemented in this study.

**Figure 2.**
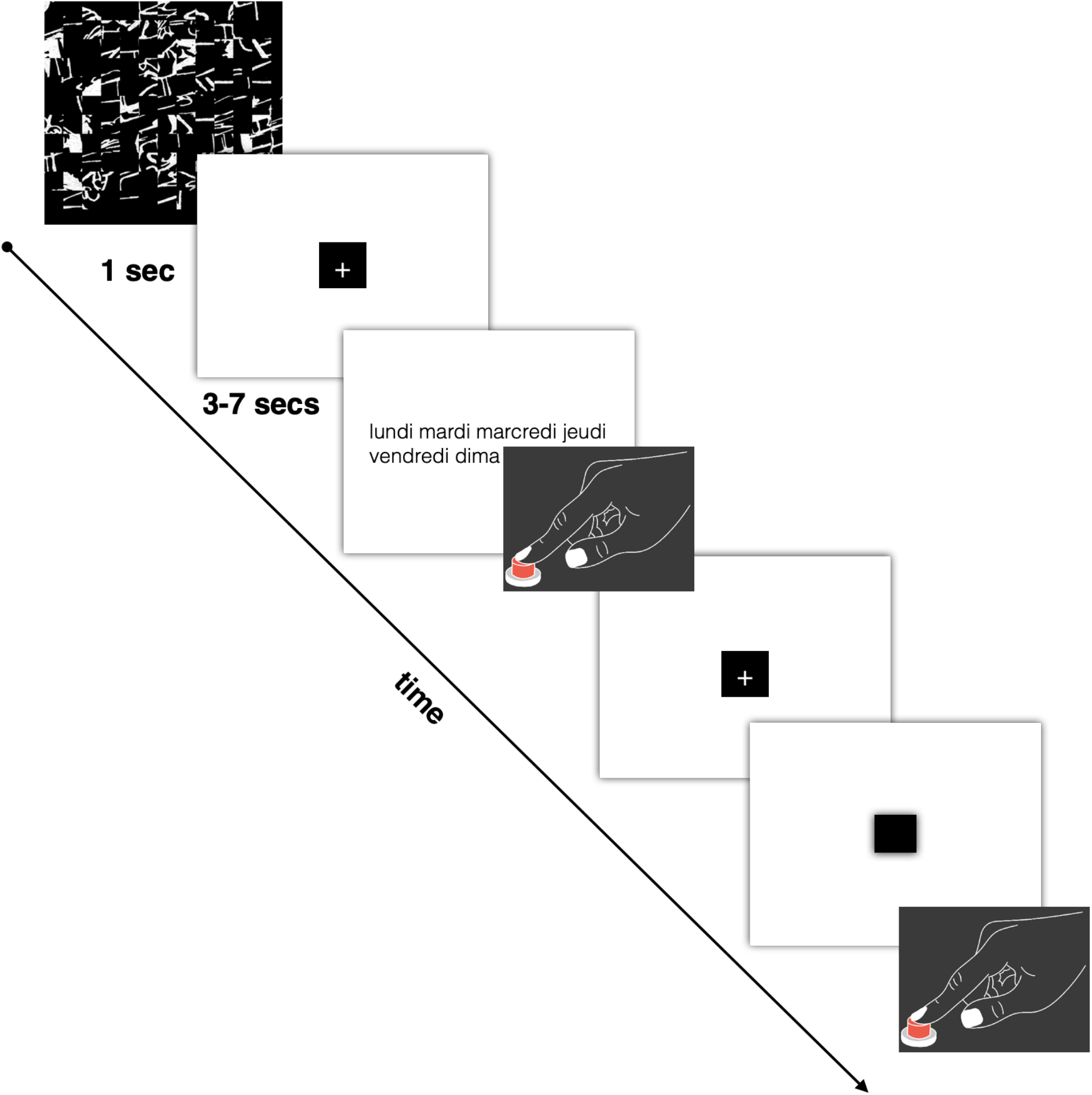
Description of the functional MRI paradigm for word-list reading. An event was initiated by presenting a scrambled picture for one second, followed by the presentation of a list of words (days, months, seasons) that participants were instructed to read. Upon task completion, participants responded with a button press. Subsequently, they had to indicate when the central cross changed into a square by clicking the button. (Figure amended from Hesling et al., 2019).

### Functional MRI

The present study included a sample of 144 healthy right-handed volunteers (72 women, 27±6 years, age range 19–53 years) from the BIL&GIN database (www.gin.cnrs.fr/en/current-research/axis2/bilgin-en/), which is a multimodal database harbouring imaging, psychometric, and genetic data to study the neural correlates of brain lateralisation (Mazoyer et al., 2016). All participants were free of diagnosed disorders and presented without a history of developmental, neurological, or psychiatric diagnoses. They were not on any medication and showed no abnormality on their structural brain MRI. All participants were French native speakers (Mazoyer et al., 2016). On average, participants spent four years at a university. All participants (n=144) were right-handed, and the group was balanced by sex (72 women).

We used the functional MRI data (fMRI) previously published in (Hesling et al., 2019) to generate t-maps of the significant voxels of functional activations during the word-list reading task.

This study concerning word reading is part of a larger study whose runs alternated this word reading task with a sentence reading task (Labache et al., 2019).

The stimuli were lists of over-learnt words, making it possible to decrease the load of lexico-semantic and syntactic processing. The lists included the days of the week, the months of the year, and the seasons. This task consisted of 13 trials lasting 14 seconds. Response times were recorded during the fMRI experiment.

The fMRI experiment included further tasks not analysed in this study. For each run, the participants were shown a scrambled drawing for 1 second, immediately followed by a central fixation crosshair. Then the participants performed a reading Word-List task and had to click on the response pad after task completion. This was followed by a low-level control task in which participants fixated on the central cross and pressed the response pad when the cross changed to a square (both stimuli covered a 0.8° × 0.8° visual area). This part of the trial lasted at least half the total trial duration. It was aimed at refocusing the participant’s attention to a non-verbal stimulus and controlling for manual motor response activation and lasted 6 - 10 seconds.

For computational purposes, t-maps of functional activations during the reading task were further characterised by measuring the average level of activations in subcortical areas (Catani and Thiebaut de Schotten, 2012) as well as in areas derived from a multimodal atlas of the brain surface (Glasser et al., 2016).

### T-maps functional MRI activations

The acquisition parameters and the pre-processing steps have been fully described (Hesling & al. 2019). Briefly, the data were acquired on a 3T Philips Intera Achieva scanner. After normalisation to the BIL&GIN template in the MNI space, the T2*-weighted echo-planar images (EPI) (TR = 2 s; TE = 35 ms; flip angle = 80°; 31 axial slices; field of view, 240 × 240 mm2; isotropic voxel size, 3.75 × 3.75 × 3.75 mm3) were initially smoothed using a 6-mm full-width at half-maximum Gaussian filter to each functional volume (194 T2*-weighted volumes). The data were processed using global linear modelling using SPM12 (https://www.fil.ion.ucl.ac.uk/spm/). For each participant, the BOLD variations of the Word-List reading task were contrasted against the cross-change detection task.

The average brain activation map of BOLD signal variations (at the voxel level) during the reading task was calculated. This activation map corresponded to the t-values extracted from the Student’s t-test for each voxel across the 144 participants. Finally, the resulting average brain map was thresholded to conserve only the significant and positive t-values (Bonferroni corrected: a voxel is significantly activated if its p-value is <0.05/n, with n = 188,561 being the number of grey matter voxels for which there is a signal across all participants).

### The disconnectome components

The Disconnectome component corresponds to the 46 principal components explaining > 99% of the variance of the white matter disconnection derived from 1333 patients with stroke lesions (see (Thiebaut de Schotten et al., 2020) for more details). Lesion data were derived from 1333 patients admitted to University College London Hospitals (UCLH) between 2001 and 2014. The study was approved by the West London & GTAC Research Ethics Committee. The data have been previously published (Thiebaut de Schotten et al., 2020; Xu et al., 2018). The lesions used in this research were non-linearly registered to the Montreal neurological institute space (2 × 2 × 2 mm, MNI; http://www.bic.mni.mcgill.ca) to allow direct individual comparisons. Notably, a second set of disconnectome components exist and can be used to replicate subsequent analyses. The disconnectome components exist as a string of values corresponding to the average level of disconnection in subcortical areas (Catani and Thiebaut de Schotten, 2012) as well as in areas derived from a multimodal atlas of the brain surface (Glasser et al., 2016) as well as neuroanatomical maps in the MNI152. Two sets of component maps — original and replication - have been provided for replication convenience of any results derived from the disconnectome (see Data availability).

### Lesion identification

1333 ischemic lesions were provided by the authors (NP) and were manually delineated on T1-weighted images. Lesion masks are derived by an unsupervised patch-based lesion-segmentation algorithm followed by manual curation as described in Xu et al. The process yields binary masks non-linearly registered to SPM’s tissue probability map (see (Xu et al., 2018) for details).

### Decoding the functional activations with the disconnectome component maps

Linear regression was applied to identify the statistical contribution of each component map (i.e. independent variables) to real group-level fMRI maps for reading (i.e. the dependent variable) (see Figure 1) in SPSS 22 (https://www.ibm.com/products/spss-statistics). The group-level fMRI maps were derived from a reading task in 144 healthy right-handed participants previously collected (Hesling et al., 2019). The 46 disconnectome components (Thiebaut de Schotten et al., 2020) explained 50% of the variance of reading activation. The linear component provided a loading (Figure 3B) for each component that was subsequently used as an independent variable in a permuted linear regression (https://fsl.fmrib.ox.ac.uk/fsl/fslwiki/Randomise). Randomise produced a map of the white matter significantly contributing to group-level reading fMRI maps.

**Figure 3.**
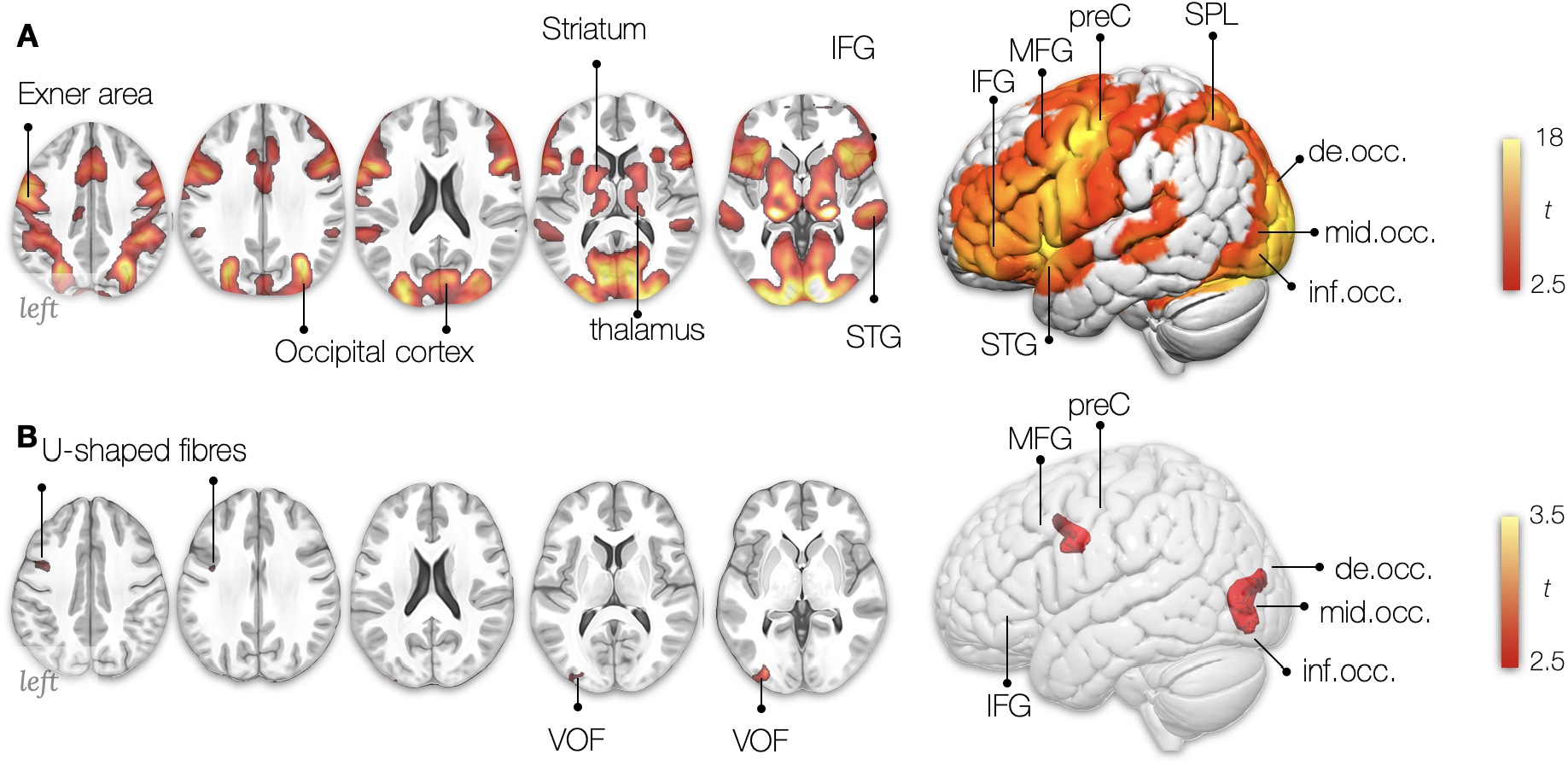
Multidimensionality of the reading network. **A** Average bilateral activation during the reading task (fMRI). **B** Locations where Disconnection components explained activations during the reading task. *t: The t-value indicates (A) the size of the difference relative to the variation in the condition and (B) the strength of the relationship between loadings (i*.*e. extracted from the linear regression between components and activation patterns) and each voxel of the components maps. In both cases, the greater the magnitude of t, the greater the evidence against the null hypothesis. IFG: inferior frontal gyrus, MFG: middle frontal gyrus, preC: precentral gyrus, SPL: superior parietal lobe, STG: superior temporal gyrus, MTG: middle temporal gyrus, de*.*occ: descending occipital gyrus, mid*.*occ: middle occipital gyrus, inf*.*occ: inferior occipital gyrus, SMG: supramarginal gyrus, AG: angular gyrus, VOF: vertical occipital fasciculus*.

Since we have two sets of component maps, we replicated the results mapped onto the white matter. The replicability of this analysis was assessed using a Pearson correlation between the results and indicated excellent replicability for fMRI reading mapping onto the white matter with a score of 83%.

#### Anatomical labelling

The white matter and grey matter were labelled manually by expert anatomists (SJF/MTdS) according to the Atlas of Human Brain Connections (Catani and Thiebaut de Schotten, 2012).

## Results

The reading task activated a bilateral network of regions in the frontal, temporal, occipital lobe and subcortical areas (Figure 3A). In the frontal lobe, activations are centred on Exner’s area (ExA) - located just between the inferior frontal gyrus’ language area and the frontal eye field (saccadic eye movements) at the intersection of the middle frontal gyrus with the precentral gyrus. Previous studies have shown that words in the visual field activate a widespread neuronal network congruent with classical language areas (Price et al., 1996). Our results replicate this observation by showing activations in the inferior frontal and superior temporal gyrus (Figure 3A).

Functional activations are also evident in the medial superior frontal gyri, the supplementary motor area (SMA). The SMA has been shown to mediate motor speech control, integrate processing between the two hemispheres and interact with subcortical structures (Hertrich et al., 2016). As a reminder, participants were asked to covertly read each word at a quiet pace to avoid them from quickly extracting the global structure of the list without actually reading it. The striatum is a subcortical structure and part of the basal ganglia that has been shown to play a role in sequencing. The frontostriatal circuit may therefore contribute to linguistic sequencing in a domain-specific manner (Chan et al., 2013), and its dysregulation has been linked to reading disorders (Hancock et al., 2017). It is thus likely that their reading included subvocal articulation. The superior, middle, and inferior occipital cortices have also been activated bilaterally.

Mapping the profiles of disconnections identified reading clusters in the occipital lobe centered around the VOF (Figure 3B).

Finally, a typical lesion was identified that showed that this disconnection profile could explain 33% of the variance in the reading task (Figure 4). Hence, this lesion will likely disconnect the white matter pathways ‘feeding’ into the activation pattern for reading. The typical lesion that causes a reading deficit is in the left hemisphere and affects Exner’s Area and the precentral gyrus in the frontal lobe; the angular gyrus in the inferior parietal lobe; the superior and middle temporal gyri, and the superior, middle and inferior occipital cortices (Figure 4C). According to the distribution of disconnection pattern maps in 1333 patients’ correlations with reading activation, such a lesion should be considered as rare –occurring in less than 0.75% of the stroke population– as indicated in Figure 4. This rare occurrence of “pure lesion” to the reading network may explain the rare occurrence of “pure alexia” patients in stroke clinics (Dejerine, 1895; Epelbaum et al., 2008; Gaillard et al., 2006; Rupareliya et al., 2017).

**Figure 4.**
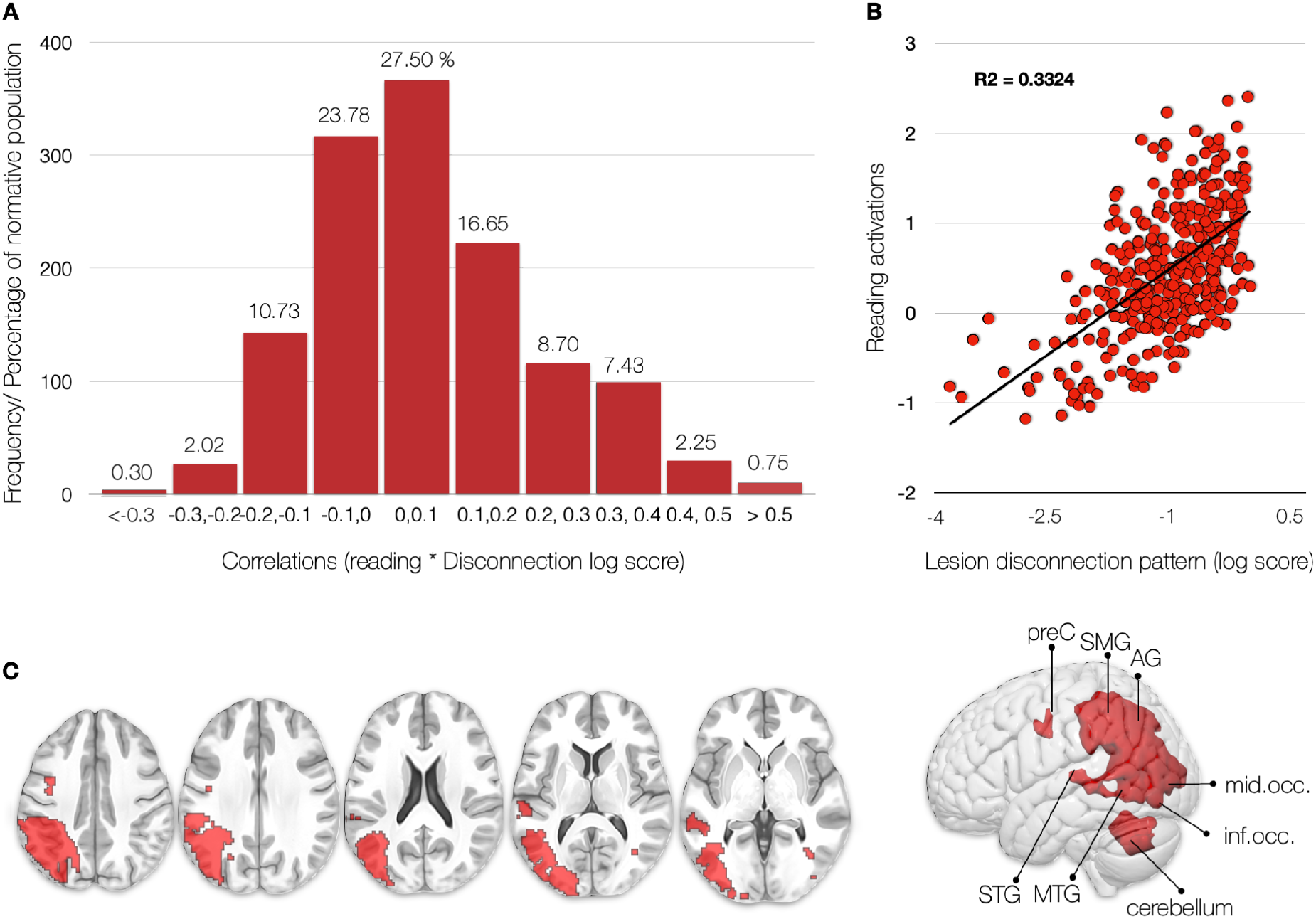
Distribution of reading activation correlations with disconnection pattern maps in 1333 patients. (A) According to our experimental framework, a “pure lesion” to the reading network would only occur in 0.75% of cases. By pure lesion, we mean lesions whose disconnection profile correlates with the reading pattern of activation with a large effect size (r > 0.5). (B) An example of a large effect size is the correlation between the pattern of functional reading task activations and the pattern of brain disconnection of a single patient, each dot corresponds to a brain region. This disconnection profile explains 33% of the variance functional reading task activations. (C) Display of the lesion pattern of the “pure lesion” to the reading network.

## Discussion

This study combined the latest research in functional MRI of reading with the disconnectome framework to revisit the neurobiology of reading. This approach couples functional MRI and structural connectivity to disentangle the circuitry, supporting a reading network in healthy controls. We propose a new, more comprehensive model of the reading network based on the convergence of functional, structural, and lesion data. First, we confirmed the functional areas activated by reading tasks during fMRI tasks. Second, we identified the critical white matter network supporting reading. Thirdly, we revealed the typical brain areas essential for reading deficits if damaged by a lesion that disconnects a reading circuitry, which would involve phylogenetically recent white matter tracts.

The cortical networks for reading have been studied for over 30 years using functional MRI (Carreiras et al., 2009; Hesling et al., 2019; Labache et al., 2019; McCrory et al., 2005; Price, 2012, 2000; Price et al., 1999; Price and Mechelli, 2005). Multiple cortical and subcortical areas have been identified as relevant for reading processes. For example, the pre-SMA and SMA-proper were associated with sequencing abstract motor plans and motor output control, respectively (Alario et al., 2006; Tremblay and Gracco, 2006). The subcortical structures, for example, the thalamus, may be involved in lexical information manipulation processes (Llano, 2013). The striatum is a subcortical structure and part of the basal ganglia associated with phonological processing and implicit sequence learning relevant to early language reading acquisition. Hyperactivation in this network has been linked to reading disorders (Hancock et al., 2017). As identified by our analysis, the network of cortical areas involved in reading matches the reading literature and is in line with the dual route theory of reading (Jobard et al., 2003). The areas of highest activation were centred in the posterior frontal lobe, anterior insula and the occipital lobes in both hemispheres (Figure 3A).

Reading familiar words, such as the days of the week and months of the year, would not necessitate sublexical assembly as the process is facilitated by lexical knowledge of the whole word. It does, however, rely on the conversion of orthographic to articulatory codes (i.e. spelling to sound conversion). As such, activations in the occipital cortex (visual), the superior temporal gyrus (auditory), and the frontal network of the IFG, MFG, preC and SMA (articulation) are very plausible (Ekert et al., 2021; Kaestner et al., 2021). fMRI does highlight all areas involved in a task by providing correlational evidence for structural-functional relationships (Damoiseaux and Greicius, 2009). Recent non-invasive and invasive brain stimulation methods allow disentangling the critical cortical regions by drawing causal inferences. Using such an approach, the precentral gyrus, which is a strongly activated hub in our fMRI analysis, is relevant for silent reading (Kaestner et al., 2021). A recent review consolidated the literature and summarised the contributions of the individual areas into a dual stream model where pseudoword reading was propagated along a dorsal parietal-frontal route while reading familiar words was processed along a ventral occipito-temporal-frontal route (Turker and Hartwigsen, 2021). In this model, the frontal areas are relevant for sound and sequencing, parietal regions for grapheme-phoneme conversion and word meaning, and the temporal areas for phonological analyses match the fMRI result we presented. A recent study identified a reading subnetwork using network-based lesion-symptom mapping that included a ventral stream with hubs in occipital, temporal, parietal and frontal areas (Li et al., 2021). The difference in the functional tasks may partially explain the different anatomy identified in this study. Li and colleagues used an overt reading task of semantically loaded content (e.g. famous people, animals). By contrast, in the task from Hesling et al., no semantic content was probed, and as such, the fMRI activations lack the classical cortical semantic hubs and circuits (Binder et al., 2009).

Within the large cortical network for reading, another area that stands out is Exner’s Area. First postulated as a neuroanatomical ‘writing centre’ by Exner (Exner, 1881), the posterior part of the middle frontal gyrus (MFG), also known as ‘Exner’s Area’, is one of the regions involved in the writing process that is highlighted in the literature (Anderson et al., 1990; Exner, 1881; Lubrano et al., 2004; Planton et al., 2013; Roux et al., 2009). Despite significant advancements in neuroimaging techniques, few studies describe patients with isolated lesions in ‘Exner’s Area’ or look at the underlying white matter connections involved in reading and writing processes. The small number of patients in the original description, in tandem with the absence of pure agraphia symptoms plus the interindividual variability in the lesion anatomy, is partially why Exner’s hypothesis is still debated (Roux et al., 2010). There are isolated reports of Exner’s area involved in reading and writing (Roux et al., 2009) and word learning in the blind (Mizuochi-Endo et al., 2021). In our study, Exner’s area has been identified as a contributing cortical region in the fMRI analysis, its disconnection from the inferior frontal gyrus and the precentral gyrus has been shown as a critical white matter connection in the disconnectome analysis, and the typical lesion that would explain most the variance in reading has also been identified in the triangle between the IFG/MFG and preC (Figure 3A,B; 4C).

Accordingly, the disconnectome analysis revealed connections to Exner’s area and Wernicke’s VOF as essential white matter circuits for reading. The frontal U-shaped fibres of the dorsal precentral gyrus connect the motor cortex and the middle frontal gyrus near Exner’s area (Catani et al., 2017) and will be referred to as ‘Exner’s U-shaped’ fibres. The VOF connects the dorsolateral and ventrolateral visual cortex (Takemura et al., 2017; Vergani et al., 2014; Yeatman et al., 2014), including the Visual Word Form Area (VWFA), a part of the ventral occipitotemporal cortex specialised in processing visual formation of words and reading (Dehaene et al., 2002; Wandell et al., 2012). A ventral temporal-occipital disconnection by lesions or surgical interventions has been linked to pure alexia (Epelbaum et al., 2008; Ng et al., 2021; Starrfelt et al., 2010). The anatomy of the VOF has been verified using tractography (Briggs et al., 2018; Keser et al., 2016; Panesar et al., 2019; Schurr et al., 2019; Yeatman et al., 2014), post-mortem dissections (Güngör et al., 2017; Palejwala et al., 2020; Vergani et al., 2014), and comparative anatomy (Takemura et al., 2017). These descriptions were scrutinised against historical post-mortem descriptions from Wernicke (Yeatman et al., 2014), Sachs (Forkel et al., 2015; Vergani et al., 2014) and the Dejerines (Bugain et al., 2021). Anatomically, the vertical occipital fibre system is distinct from the parietal-temporal branch of the arcuate fasciculus (e.g. posterior segment) in its trajectories and cortical terminations. The intralobar vertical occipital fasciculus projects between the ventral and dorsal occipital cortices while the interlobar posterior segment of the arcuate fasciculus connects posterior temporal and inferior parietal regions (i.e. supramarginal and angular gyrus) (Weiner et al., 2017). A recent review identified an association between the VOF and visual tasks, including object recognition and integrating the dorsal and ventral stream during hand actions (Budisavljevic et al., 2018; Forkel et al., 2021). The VOF terminations have also been shown to predict reading-related responses in the human ventral temporal cortex (Grotheer et al., 2021) and early literacy skills in children (Broce et al., 2019). In the clinical setting, the VOF has recently also been associated with visual scores in assessments of patients with multiple sclerosis (Abdolalizadeh et al., 2022). Hence, the disconnectome identification of U-shaped fibres and the VOF as critical connections supporting literacy is a plausible explanation, although other U-shaped fibres, yet unidentified, may also overlap with this result.

These U-shaped fibres might have evolved later on in the phylogenetic tree. The emergence of literacy necessitated accessing our knowledge of spoken language through a novel modality, one that was never anticipated by evolution: vision. While we as a species have been using representations of visual objects and spoken sounds using visual signs (e.g. hieroglyphs) for a long time, our brains mostly evolved for millions of years in a world where literacy, or any form thereof, did not exist (Hodgson, 2019). It is argued that reading developed some 5000 years ago, a time frame too short for significant evolutionary changes. As a consequence, some say that reading would rely on brain regions that initially evolved to process vision (e.g. VOF) and spoken language (e.g. Exner’s U-shaped fibres) (Dehaene et al., 2010) and the connections at their intersection (Yeatman and White, 2021). However, the specificity of the VWFA has been challenged (Price and Devlin, 2004, 2003). Some might argue that the reading brain does not exist specifically but emerged through brain plasticity in the grey and the white matter (Carreiras et al., 2009; Dehaene et al., 2010; Thiebaut de Schotten et al., 2014).

This study identified the critical role of fronto-parietal U-shaped fibres underneath Exner’s area. The superficial white matter, which includes short-ranged intergyral U-shaped fibres, is more variable between people and a more recent addition to brain evolution (Croxson et al., 2018). If we consider the emergence of literacy as a late appearance in the evolution of the brain, it would be no surprise that reading relies on a more adaptable and flexible white matter network, i.e. U-shaped fibres. This is because there is a gradient of variability where the more superficial white matter is more variable while the deeper white matter is more conserved across individuals. The U-shaped fibres are the most superficial connections in the human brain. They might consequently adjust to new demands as the most variable areas are associated with higher cognitive functions in humans (Croxson et al., 2018). The VOF is also a short-ranging intracortical connection and, as such, is located superficially. The critical involvement of U-shaped fibres and short-range connections in reading may reflect a recent adaptation to the advancement of human skills.

We have demonstrated in this study that the most likely lesion to disrupt the reading circuitry was located in the inferior parietal lobe, the posterior, superior, and middle temporal cortex, the superior, middle and inferior occipital cortex, and Exner’s area of the left hemisphere (Figure 4). The typical lesion affected the right hemisphere’s white matter behind the inferior temporal gyrus. This lesion would consist of the language areas identified from the fMRI task, the U-shaped fibres, and the vertical occipital fasciculus, unifying the results determined by functional imaging and disconnectome mapping. This result is in line with recent neuroimaging studies suggesting that learned orthographic knowledge is associated with the left inferior temporal-occipital cortex and fusiform gyrus (Cohen and Dehaene, 2004). As well as, the literature suggests that lesions to that area should lead to reading impairments. For example, a voxel-based lesion-symptom study in 111 stroke patients identified the critical lesion areas for reading (left occipitotemporal cortex, and inferior parietal cortex), writing (left inferior parietal lobe, superior temporal cortex including Heschl’s gyrus, and sensorimotor cortex), single-word reading (left temporo-occipital cortex, posterior lateral temporal, and inferior parietal cortex), and sentence reading (left temporo-occipital cortex, lateral temporal cortex, superior portion of the temporal pole, inferior and posterior insula) (Baldo et al., 2018). The typical lesion identified from our cohort of 1333 patients matches these results for single-word reading with the addition of Exner’s area and the cerebellum. The cerebellum has been discussed in the clinical dyslexia literature as the cerebellar deficit hypothesis, which suggests that cerebellar anomalies could give rise to developmental dyslexia by impairing articulatory/phonological monitoring (Nicolson et al., 2001). The discussion of the cerebellum’s role in language functions has recently gained momentum (Mariën et al., 2014; Mariën and Borgatti, 2018; Murdoch, 2010). An fMRI study in healthy adults with differing reading proficiency has also identified cerebellar activations associated with motor/articulatory processing (Cullum et al., 2019).

Decoding task-based functional MRI imaging using a disconnectome framework is a new and promising method to link structural connectivity with function and behaviour using brain disconnections. This method has already been successfully applied to various cognitive functions and behaviours (Dulyan et al., 2021; Thiebaut de Schotten et al., 2020) and offers an indirect measure of the structural connectivity supporting functional activations. To directly predict reading impairments and estimate measures of reading impairments, we provide the maps associated with this study on Neurovault for future research.

Nevertheless, caution should be applied to these maps as patient results might vary according to demographics, reading abilities or literacy skills. Further, we employed the reading task based on automatic stimuli and tapped into basic language processes. Accordingly, this task is not loaded linguistically and likely bypasses complex language processes. However, complex linguistic fMRI tasks would likely activate the whole brain rendering our decoding impossible. Finally, the low requirement of our reading task makes it suitable for the clinical settings opening up new opportunities for future clinical investigations.

Of all human inventions, the ingenuity of the written language has been momentous and far-reaching in human evolution. Literacy has enabled a considerable increase in the rate and efficacy of cultural evolution. The cultural evolution of society and individuals relies on knowledge transmission within and across generations, which was greatly facilitated by literacy. Neuroimaging offers the tools to uncover the brain circuitry that enables literacy, and this paper is a novel contribution to deciphering the neurobiology of reading.

Overall, we demonstrated that the reading circuitry can be decoded in healthy participants’ functional activations using a disconnectome approach and identified Exner’s U-shaped fibres and the vertical occipital fasciculus as critical white matter connections to reading.

## Acknowledgements

The authors would like to thank the Groupe d’imagerie neurofonctionnelle (GIN) whiteboard team for helpful discussions, with special thanks to Valentina Pacella and Lia Talozzi.

## Data availability

Disconnectome maps and lesions were downloaded from the supplementary material of (Thiebaut de Schotten et al., 2020) The two sets of component maps and the atlas of white matter function (original and replication) are available at https://identifiers.org/neurovault.collection:7735. The scatterplot (fMRI and disconnectome) was done using the available python code http://www.bcblab.com/BCB/Coding/Entries/2019/3/15_Produce_scatterplots_of_correlation_between_2_.nii_files.html.

Maps associated with this research are available on Neurovault.org (Collection: https://identifiers.org/neurovault.collection:12309.

Individual files: fMRI: https://identifiers.org/neurovault.image:777434; Disco map: https://identifiers.org/neurovault.image:777435; typical lesion: https://identifiers.org/neurovault.image:777436).

## Bibliography

Abdolalizadeh, A., Mohammadi, S., Aarabi, M.H., 2022. The forgotten tract of vision in multiple sclerosis: vertical occipital fasciculus, its fiber properties, and visuospatial memory. Brain Struct. Funct.

Alario, F.-X., Chainay, H., Lehericy, S., Cohen, L., 2006. The role of the supplementary motor area (SMA) in word production. Brain Res. 1076, 129–143.

Anderson, S.W., Damasio, A.R., Damasio, H., 1990. Troubled letters but not numbers. Domain specific cognitive impairments following focal damage in frontal cortex. Brain 113 (Pt 3), 749–766.

Baldo, J.V., Kacinik, N., Ludy, C., Paulraj, S., Moncrief, A., Piai, V., Curran, B., Turken, A., Herron, T., Dronkers, N.F., 2018. Voxel-based lesion analysis of brain regions underlying reading and writing. Neuropsychologia 115, 51–59.

Binder, J.R., Desai, R.H., Graves, W.W., Conant, L.L., 2009. Where is the semantic system? A critical review and meta-analysis of 120 functional neuroimaging studies. Cereb. Cortex 19, 2767–2796.

Briggs, R.G., Conner, A.K., Sali, G., Rahimi, M., Baker, C.M., Burks, J.D., Glenn, C.A., Battiste, J.D., Sughrue, M.E., 2018. A Connectomic Atlas of the Human Cerebrum-Chapter 16: Tractographic Description of the Vertical Occipital Fasciculus. Oper Neurosurg (Hagerstown) 15, S456–S461.

Broce, I.J., Bernal, B., Altman, N., Bradley, C., Baez, N., Cabrera, L., Hernandez, G., De Feria, A., Dick, A.S., 2019. Fiber pathways supporting early literacy development in 5-8-year-old children. Brain Cogn. 134, 80–89.

Buchsbaum, B.R., Olsen, R.K., Koch, P.F., Kohn, P., Kippenhan, J.S., Berman, K.F., 2005. Reading, hearing, and the planum temporale. Neuroimage 24, 444–454.

Budisavljevic, S., Dell’Acqua, F., Castiello, U., 2018. Cross-talk connections underlying dorsal and ventral stream integration during hand actions. Cortex 103, 224–239.

Bugain, M., Dimech, Y., Torzhenskaya, N., Thiebaut de Schotten, M., Caspers, S., Muscat, R., Bajada, C.J., 2021. Occipital Intralobar fasciculi: a description, through tractography, of three forgotten tracts. Commun Biol 4, 433.

Burton, M.W., Small, S.L., Blumstein, S.E., 2000. The role of segmentation in phonological processing: an fMRI investigation. J. Cogn. Neurosci. 12, 679–690.

Carreiras, M., Seghier, M.L., Baquero, S., Estévez, A., Lozano, A., Devlin, J.T., Price, C.J., 2009. An anatomical signature for literacy. Nature 461, 983–986.

Catani, M., Jones, D.K., Donato, R., Ffytche, D.H., 2003. Occipito-temporal connections in the human brain. Brain 126, 2093–2107.

Catani, M., Robertsson, N., Beyh, A., Huynh, V., de Santiago Requejo, F., Howells, H., Barrett, R.L.C., Aiello, M., Cavaliere, C., Dyrby, T.B., Krug, K., Ptito, M., D’Arceuil, H., Forkel, S.J., Dell’Acqua, F., 2017. Short parietal lobe connections of the human and monkey brain. Cortex 97, 339–357.

Catani, M., Thiebaut de Schotten, M., 2012. Atlas of Human Brain Connections.

Chan, S.-H., Ryan, L., Bever, T.G., 2013. Role of the striatum in language: Syntactic and conceptual sequencing. Brain Lang. 125, 283–294.

Cohen, L., Dehaene, S., 2004. Specialization within the ventral stream: the case for the visual word form area. Neuroimage 22, 466–476.

Cohen, L., Dehaene, S., Naccache, L., Lehéricy, S., Dehaene-Lambertz, G., Hénaff, M.A., Michel, F., 2000. The visual word form area: spatial and temporal characterization of an initial stage of reading in normal subjects and posterior split-brain patients. Brain 123 (Pt 2), 291–307.

Cohen, L., Henry, C., Dehaene, S., Martinaud, O., Lehéricy, S., Lemer, C., Ferrieux, S., 2004. The pathophysiology of letter-by-letter reading. Neuropsychologia 42, 1768–1780.

Croxson, P.L., Forkel, S.J., Cerliani, L., Thiebaut de Schotten, M., 2018. Structural Variability Across the Primate Brain: A Cross-Species Comparison. Cereb. Cortex 28, 3829–3841.

Cullum, A., Hodgetts, W.E., Milburn, T.F., Cummine, J., 2019. Cerebellar Activation During Reading Tasks: Exploring the Dichotomy Between Motor vs. Language Functions in Adults of Varying Reading Proficiency. Cerebellum 18, 688–704.

Damoiseaux, J.S., Greicius, M.D., 2009. Greater than the sum of its parts: a review of studies combining structural connectivity and resting-state functional connectivity. Brain Struct. Funct. 213, 525–533.

Dehaene, S., Cohen, L., 2011. The unique role of the visual word form area in reading. Trends Cogn. Sci. 15, 254–262.

Dehaene, S., Le Clec’H, G., Poline, J.-B., Le Bihan, D., Cohen, L., 2002. The visual word form area: a prelexical representation of visual words in the fusiform gyrus. Neuroreport 13, 321–325.

Dehaene, S., Pegado, F., Braga, L.W., Ventura, P., Nunes Filho, G., Jobert, A., Dehaene-Lambertz, G., Kolinsky, R., Morais, J., Cohen, L., 2010. How learning to read changes the cortical networks for vision and language. Science 330, 1359–1364.

Dejerine, J., 1895. Anatomie des Centres Nerveux. Rueff et Cie, Paris.

Dulyan, L., Talozzi, L., Pacella, V., Corbetta, M., Forkel, S.J., de Schotten, M.T., 2021. Longitudinal prediction of motor dysfunction after stroke: a disconnectome study. bioRxiv.

Ekert, J.O., Lorca-Puls, D.L., Gajardo-Vidal, A., Crinion, J.T., Hope, T.M.H., Green, D.W., Price, C.J., 2021. A functional dissociation of the left frontal regions that contribute to single word production tasks. Neuroimage 245, 118734.

Epelbaum, S., Pinel, P., Gaillard, R., Delmaire, C., Perrin, M., Dupont, S., Dehaene, S., Cohen, L., 2008. Pure alexia as a disconnection syndrome: new diffusion imaging evidence for an old concept. Cortex 44, 962–974.

Exner, S., 1881. Untersuchungen über die Lokalisation der Functionen in der Grosshirnrinde des Menschen … von Prof. Sigmund Exner. W. Braumüller.

Fiset, D., Gosselin, F., Blais, C., Arguin, M., 2006. Inducing letter-by-letter dyslexia in normal readers. J. Cogn. Neurosci. 18, 1466–1476.

Flinker, A., Korzeniewska, A., Shestyuk, A.Y., Franaszczuk, P.J., Dronkers, N.F., Knight, R.T., Crone, N.E., 2015. Redefining the role of Broca’s area in speech. Proc. Natl. Acad. Sci. U. S. A. 112, 2871–2875.

Forkel, S.J., Friedrich, P., Thiebaut de Schotten, M., Howells, H., 2021. White matter variability, cognition, and disorders: a systematic review. Brain Struct. Funct.

Forkel, S.J., Mahmood, S., Vergani, F., Catani, M., 2015. The white matter of the human cerebrum: part I The occipital lobe by Heinrich Sachs. Cortex 62, 182–202.

Forkel, S.J., Thiebaut de Schotten, M., Kawadler, J.M., Dell’Acqua, F., Danek, A., Catani, M., 2014. The anatomy of fronto-occipital connections from early blunt dissections to contemporary tractography. Cortex 56, 73–84.

Friederici, A.D., 2002. Towards a neural basis of auditory sentence processing. Trends Cogn. Sci. 6, 78–84.

Gaillard, R., Naccache, L., Pinel, P., Clémenceau, S., Volle, E., Hasboun, D., Dupont, S., Baulac, M., Dehaene, S., Adam, C., Cohen, L., 2006. Direct intracranial, FMRI, and lesion evidence for the causal role of left inferotemporal cortex in reading. Neuron 50, 191–204.

Glasser, M.F., Coalson, T.S., Robinson, E.C., Hacker, C.D., Harwell, J., Yacoub, E., Ugurbil, K., Andersson, J., Beckmann, C.F., Jenkinson, M., Smith, S.M., Van Essen, D.C., 2016. A multi-modal parcellation of human cerebral cortex. Nature 536, 171–178.

Grotheer, M., Yeatman, J., Grill-Spector, K., 2021. White matter fascicles and cortical microstructure predict reading-related responses in human ventral temporal cortex. Neuroimage 227, 117669.

Güngör, A., Baydin, S., Middlebrooks, E.H., Tanriover, N., Isler, C., Rhoton, A.L., Jr, 2017. The white matter tracts of the cerebrum in ventricular surgery and hydrocephalus. J. Neurosurg. 126, 945–971.

Hagoort, P., 2005. On Broca, brain, and binding: a new framework. Trends Cogn. Sci. 9, 416–423.

Hancock, R., Richlan, F., Hoeft, F., 2017. Possible roles for fronto-striatal circuits in reading disorder. Neurosci. Biobehav. Rev. 72, 243–260.

Hertrich, I., Dietrich, S., Ackermann, H., 2016. The role of the supplementary motor area for speech and language processing. Neurosci. Biobehav. Rev. 68, 602–610.

Hesling, I., Labache, L., Joliot, M., Tzourio-Mazoyer, N., 2019. Large-scale plurimodal networks common to listening to, producing and reading word lists: an fMRI study combining task-induced activation and intrinsic connectivity in 144 right-handers. Brain Struct. Funct. 224, 3075–3094.

Hodgson, D., 2019. The origin, significance, and development of the earliest geometric patterns in the archaeological record. Journal of Archaeological Science: Reports 24, 588–592.

Jobard, G., Crivello, F., Tzourio-Mazoyer, N., 2003. Evaluation of the dual route theory of reading: a metanalysis of 35 neuroimaging studies. Neuroimage 20, 693–712.

Kaestner, E., Thesen, T., Devinsky, O., Doyle, W., Carlson, C., Halgren, E., 2021. An Intracranial Electrophysiology Study of Visual Language Encoding: The Contribution of the Precentral Gyrus to Silent Reading. J. Cogn. Neurosci. 33, 2197–2214.

Keser, Z., Ucisik-Keser, F.E., Hasan, K.M., 2016. Quantitative Mapping of Human Brain Vertical-Occipital Fasciculus. J. Neuroimaging 26, 188–193.

Labache, L., Joliot, M., Saracco, J., Jobard, G., Hesling, I., Zago, L., Mellet, E., Petit, L., Crivello, F., Mazoyer, B., Tzourio-Mazoyer, N., 2019. A SENtence Supramodal Areas AtlaS (SENSAAS) based on multiple task-induced activation mapping and graph analysis of intrinsic connectivity in 144 healthy right-handers. Brain Struct. Funct. 224, 859–882.

Li, M., Song, L., Zhang, Y., Han, Z., 2021. White matter network of oral word reading identified by network-based lesion-symptom mapping. iScience 24, 102862.

Llano, D.A., 2013. Functional imaging of the thalamus in language. Brain Lang. 126, 62–72.

López-Barroso, D., Catani, M., Ripollés, P., Dell’Acqua, F., Rodríguez-Fornells, A., de Diego-Balaguer, R., 2013. Word learning is mediated by the left arcuate fasciculus. Proc. Natl. Acad. Sci. U. S. A. 110, 13168–13173.

Lubrano, V., Roux, F.-E., Démonet, J.-F., 2004. Writing-specific sites in frontal areas: a cortical stimulation study. J. Neurosurg. 101, 787–798.

Luzzatti, C., Mondini, S., Semenza, C., 2001. Lexical representation and processing of morphologically complex words: evidence from the reading performance of an Italian agrammatic patient. Brain Lang. 79, 345–359.

Mariën, P., Ackermann, H., Adamaszek, M., Barwood, C.H.S., Beaton, A., Desmond, J., De Witte, E., Fawcett, A.J., Hertrich, I., Küper, M., Leggio, M., Marvel, C., Molinari, M., Murdoch, B.E., Nicolson, R.I., Schmahmann, J.D., Stoodley, C.J., Thürling, M., Timmann, D., Wouters, E., Ziegler, W., 2014. Consensus paper: Language and the cerebellum: an ongoing enigma. Cerebellum 13, 386–410.

Mariën, P., Borgatti, R., 2018. Language and the cerebellum. Handb. Clin. Neurol. 154, 181–202.

Mazoyer, B., Mellet, E., Perchey, G., Zago, L., Crivello, F., Jobard, G., Delcroix, N., Vigneau, M., Leroux, G., Petit, L., Joliot, M., Tzourio-Mazoyer, N., 2016. BIL&GIN: A neuroimaging, cognitive, behavioral, and genetic database for the study of human brain lateralization. Neuroimage 124, 1225–1231.

McCrory, E.J., Mechelli, A., Frith, U., Price, C.J., 2005. More than words: a common neural basis for reading and naming deficits in developmental dyslexia? Brain 128, 261–267.

Mizuochi-Endo, T., Itou, K., Makuuchi, M., Kato, B., Ikeda, K., Nakamura, K., 2021. Graphomotor memory in Exner’s area enhances word learning in the blind. Communications Biology.

Murdoch, B.E., 2010. The cerebellum and language: historical perspective and review. Cortex 46, 858–868.

Murphy, K.A., Jogia, J., Talcott, J.B., 2019. On the neural basis of word reading: A meta-analysis of fMRI evidence using activation likelihood estimation. J. Neurolinguistics 49, 71–83.

Ng, S., Moritz-Gasser, S., Lemaitre, A.-L., Duffau, H., Herbet, G., 2021. White matter disconnectivity fingerprints causally linked to dissociated forms of alexia. Commun Biol 4, 1413.

Nicolson, R.I., Fawcett, A.J., Dean, P., 2001. Developmental dyslexia: the cerebellar deficit hypothesis. Trends Neurosci. 24, 508–511.

Palejwala, A.H., O’Connor, K.P., Pelargos, P., Briggs, R.G., Milton, C.K., Conner, A.K., Milligan, T.M., O’Donoghue, D.L., Glenn, C.A., Sughrue, M.E., 2020. Anatomy and white matter connections of the lateral occipital cortex. Surg. Radiol. Anat. 42, 315–328.

Panesar, S.S., Belo, J.T.A., Yeh, F.-C., Fernandez-Miranda, J.C., 2019. Structure, asymmetry, and connectivity of the human temporo-parietal aslant and vertical occipital fasciculi. Brain Struct. Funct. 224, 907–923.

Patterson, K., Hodges, J.R., 1992. Deterioration of word meaning: Implications for reading. Neuropsychologia.

Philipose, L.E., Gottesman, R.F., Newhart, M., Kleinman, J.T., Herskovits, E.H., Pawlak, M.A., Marsh, E.B., Davis, C., Heidler-Gary, J., Hillis, A.E., 2007. Neural regions essential for reading and spelling of words and pseudowords. Ann. Neurol. 62, 481–492.

Planton, S., Jucla, M., Roux, F.-E., Démonet, J.-F., 2013. The “handwriting brain”: A meta-analysis of neuroimaging studies of motor versus orthographic processes. Cortex 49, 2772–2787.

Price, C.J., 2000. The anatomy of language: contributions from functional neuroimaging. J. Anat. 197 Pt 3, 335–359.

Price, C.J., 2012. A review and synthesis of the first 20 years of PET and fMRI studies of heard speech, spoken language and reading. Neuroimage 62, 816–847.

Price, C.J., Devlin, J.T., 2003. The myth of the visual word form area. Neuroimage 19, 473–481.

Price, C.J., Devlin, J.T., 2004. The pro and cons of labelling a left occipitotemporal region: “the visual word form area.” Neuroimage.

Price, C.J., Green, D.W., von Studnitz, R., 1999. A functional imaging study of translation and language switching. Brain 122 (Pt 12), 2221–2235.

Price, C.J., Mechelli, A., 2005. Reading and reading disturbance. Curr. Opin. Neurobiol. 15, 231–238.

Price, C.J., Wise, R.J., Frackowiak, R.S., 1996. Demonstrating the implicit processing of visually presented words and pseudowords. Cereb. Cortex 6, 62–70.

Roux, F.-E., Draper, L., Köpke, B., Démonet, J.-F., 2010. Who actually read Exner? Returning to the source of the frontal “writing centre” hypothesis. Cortex 46, 1204–1210.

Roux, F.-E., Dufor, O., Giussani, C., Wamain, Y., Draper, L., Longcamp, M., Démonet, J.-F., 2009. The graphemic/motor frontal area Exner’s area revisited. Ann. Neurol. 66, 537–545.

Rupareliya, C., Naqvi, S., Hejazi, S., 2017. Alexia Without Agraphia: A Rare Entity. Cureus.

Schurr, R., Filo, S., Mezer, A.A., 2019. Tractography delineation of the vertical occipital fasciculus using quantitative T1 mapping. Neuroimage 202, 116121.

Sihvonen, A.J., Virtala, P., Thiede, A., Laasonen, M., Kujala, T., 2021. Structural white matter connectometry of reading and dyslexia. Neuroimage 241, 118411.

Starrfelt, R., Habekost, T., Gerlach, C., 2010. Visual processing in pure alexia: a case study. Cortex 46, 242–255.

Starrfelt, R., Shallice, T., 2014. What’s in a name? The characterization of pure alexia. Cogn. Neuropsychol. 31, 367–377.

Takemura, H., Pestilli, F., Weiner, K.S., Keliris, G.A., Landi, S.M., Sliwa, J., Ye, F.Q., Barnett, M.A., Leopold, D.A., Freiwald, W.A., Logothetis, N.K., Wandell, B.A., 2017. Occipital White Matter Tracts in Human and Macaque. Cereb. Cortex 27, 3346–3359.

Thiebaut de Schotten, M., Cohen, L., Amemiya, E., Braga, L.W., Dehaene, S., 2014. Learning to read improves the structure of the arcuate fasciculus. Cereb. Cortex 24, 989–995.

Thiebaut de Schotten, M., Foulon, C., Nachev, P., 2020. Brain disconnections link structural connectivity with function and behaviour. Nat. Commun. 11, 5094.

Tremblay, P., Gracco, V.L., 2006. Contribution of the frontal lobe to externally and internally specified verbal responses: fMRI evidence. Neuroimage 33, 947–957.

Turker, S., Hartwigsen, G., 2021. Exploring the neurobiology of reading through non-invasive brain stimulation: A review. Cortex 141, 497–521.

Vanderauwera, J., De Vos, A., Forkel, S.J., Catani, M., Wouters, J., Vandermosten, M., Ghesquière, P., 2018. Neural organization of ventral white matter tracts parallels the initial steps of reading development: A DTI tractography study. Brain Lang. 183, 32–40.

Vergani, F., Mahmood, S., Morris, C.M., Mitchell, P., Forkel, S.J., 2014. Intralobar fibres of the occipital lobe: a post mortem dissection study. Cortex 56, 145–156.

Wandell, B.A., Rauschecker, A.M., Yeatman, J.D., 2012. Learning to see words. Annu. Rev. Psychol. 63, 31–53.

Weiner, K.S., Yeatman, J.D., Wandell, B.A., 2017. The posterior arcuate fasciculus and the vertical occipital fasciculus. Cortex 97, 274–276.

Xu, T., Rolf Jäger, H., Husain, M., Rees, G., Nachev, P., 2018. High-dimensional therapeutic inference in the focally damaged human brain. Brain 141, 48–54.

Yeatman, J.D., Dougherty, R.F., Ben-Shachar, M., Wandell, B.A., 2012. Development of white matter and reading skills. Proc. Natl. Acad. Sci. U. S. A. 109, E3045–53.

Yeatman, J.D., Dougherty, R.F., Rykhlevskaia, E., 2011. Anatomical properties of the arcuate fasciculus predict phonological and reading skills in children. Journal of cognitive.

Yeatman, J.D., Weiner, K.S., Pestilli, F., Rokem, A., Mezer, A., Wandell, B.A., 2014. The vertical occipital fasciculus: a century of controversy resolved by in vivo measurements. Proc. Natl. Acad. Sci. U. S. A. 111, E5214–23.

Yeatman, J.D., White, A.L., 2021. Reading: The Confluence of Vision and Language. Annu Rev Vis Sci 7, 487–517.

Dejerine, J. (1895). Anatomie des Centres Nerveux. Paris, Rueff et Cie.

